# Co-evolutionary signals from *Burkholderia pseudomallei* genomics identify its survival strategies and highlight improving environmental health as prevention policy

**DOI:** 10.1101/2020.08.11.245894

**Authors:** Claire Chewapreecha, Johan Pensar, Supaksorn Chattagul, Maiju Pesonen, Apiwat Sangphukieo, Phumrapee Boonklang, Chotima Potisap, Sirikamon Koosakulnirand, Edward J Feil, Susanna Dunachie, Narisara Chantratita, Direk Limmathurotsakul, Sharon J Peacock, Nick PJ Day, Julian Parkhill, Nicholas R Thomson, Rasana W Sermswan, Jukka Corander

## Abstract

**Background:** The soil bacterium *Burkholderia pseudomallei* is the causative agent of melioidosis. It kills up to 40% of cases and contributes to human morbidity and mortality in many tropical and sub-tropical countries. As no vaccines are currently available, prevention is the key health policy and is achieved by avoiding direct contact with soil and standing water. The pathogen notoriously persists in ranges of environmental conditions which make disease prevention difficult. We aimed to scan *B. pseudomallei* genomes for signals of evolutionary adaptations that allow it to thrive across environmental conditions, which should ultimately inform prevention policy.

**Methods:** We conducted three layers of analyses: a genome-wide epistasis and co-selection study (GWES) on 2,011 *B. pseudomallei* genomes to detect signals of co-selection; gene expression analyses across 82 diverse physical, chemical, biological and infectious conditions to identify specific conditions in which such selection might have acted; and gene knockout assays to confirm the function of the co-selection hotspot.

**Findings:** We uncovered 13,061 mutation pairs in distinct genes and non-coding RNA that have been repeatedly co-selected through *B. pseudomallei* evolution. Genes under co-selection displayed marked expression correlation when *B. pseudomallei* was subjected to physical stress conditions including temperature stress, osmotic stress, UV radiation, and nutrient deprivation; highlighting these conditions as the major evolutionary driving forces for this bacterium. We identified a putative adhesin (*BPSL1661*) as a hub of co-selection signals, experimentally confirmed the role of *BPSL1661* under nutrient deprivation, and explored the functional basis of the co-selection gene network surrounding *BPSL1661* in facilitating bacterial survival under nutrient depletion.

**Interpretation:** Our findings suggest that *B. pseudomallei* has a selective advantage to survive nutrient-limited conditions. Anthropogenic activities such as shifting cultivation systems with more frequent rotations of cropping and shortened fallow periods or continuous cultivation of cash crops could directly or indirectly contribute to loss of soil nutrient; these may lead to the preferential survival of *B. pseudomallei* and a subsequent rise of melioidosis. Successful disease control for melioidosis needs to consider improving environmental health in addition to current preventive efforts.

**Funding:** Wellcome Trust, European Research Council, UK Department of Health, Thailand Research Fund and Khon Kaen University

**Research in context:** *Evidence before this study:* We searched PubMed with terms (co-selection AND bacteria AND population) with no date or language restrictions from database inception until April 11, 2021. We identified 44 publications of which four were conducted at a genome-wide scale. These four studies were performed on human-restricted pathogens, detected co-selection of antibiotic resistance gene networks which highlight the use of antibiotics as major selection pressures and further inform treatment options. However, none of these studies were performed on *Burkholderia pseudomallei* or other opportunistic pathogens that have been adapted to both natural and host environments. The selection pressures exerted on these pathogens and the genetic determinants allowed for their adaptations remain unclear, which limit our understanding on the bacterial biology and the information used for disease control.

*Added value of this study:* Based on genomes of 2,011 *B. pseudomallei* collected from melioidosis endemic areas, we identified and confirmed genetic signals for co-selection. Using transcriptome profiling covering a broad spectrum of conditions and exposures, we showed that genes under co-selection displayed marked expression correlation under physical stress conditions with the gene at the co-selection hotspot conditionally expressed under nutrient starvation. Furthermore, we experimentally validated the function of the hotspot gene and demonstrated that unlike host-restricted pathogens, the *B. pseudomallei* co-selection network does not facilitate host infection but is focused on bacterial survival in a harsh environment, particularly under nutrient depletion. Aside from providing a data resource, the study also showcases the power of combined genetics, transcriptomics and functional analysis as a tool for biology discovery.

*Implications of all available evidence:* Our findings provide evolutionary and biological evidence for preferential survival of *B. pseudomallei* under nutrient starvation. Agricultural practice that induces soil loss, which is not uncommon in melioidosis endemic areas has been linked to soil nutrient depletion and may contribute to the prevalence of *B. pseudomallei* and a consequent rise of melioidosis in these regions. Successful melioidosis control has to consider environmental health in addition to existing prevention policy.

## Introduction

The environmental bacterium *Burkholderia pseudomallei* has been increasingly recognised as an emerging human and animal pathogen and a cause of melioidosis, a rapidly fatal infectious disease that affects and kills an estimated number of 165,000 and 89,000 patients globally per year.^1^ Melioidosis results from inoculation, ingestion or inhalation of *B. pseudomallei* from contaminated soil, dust, surface water or water droplets.^2^ Currently, vaccines are not available. The key health policy is thus based on avoiding direct contact with the contaminated environment and individuals with unavoidable contact, such as rice farmers, wearing protective clothing or footwear. The bacterium can be isolated from the soil in many tropical and sub-tropical regions. It is most abundant at depths ≥ 10 cm from the surface but can move from deeper soil layers to the soil surface during the rainy season and multiply; this has been linked to an increase in disease incidence after heavy rainfalls^2^. *B. pseudomallei* can survive extreme environmental conditions ranging from dry terrain in deserts^3^ to distilled water (no nutrients)^4^; the latter ongoing experiment has been run for over 25 years. Our previous work links the presence of *B. pseudomallei* with nutrient-depleted soil and soil modified by long-term human activity; thereby connecting human manipulation of soil physicochemistry that promotes the abundance of this species.^5^ The bacterium’s ability to persist in diverse and tough environmental conditions has challenged disease prevention efforts. The identification of the genetic factors that mediate bacterial adaptations to ranges of environmental conditions has been a long-standing research objective, yet has been little explored at a genome-wide or a population scale. In this study, we aimed to determine the genetic loci that have been repeatedly selected through *B. pseudomallei* evolution and link these loci to environmental conditions that the species has adapted to, which ultimately helps inform prevention policy.

## Methods

### Study design

We conducted three complementary tests: a co-selection analysis to scan for any SNP-SNP pairs that are mutually detected more frequently than expected as a result of selection pressures; a condition-wide transcriptome analysis to identify conditions under which co-selected gene-gene pairs are up-regulated; and a knockout assay of the co-selection hotspot gene to validate the gene function.

### Whole-genome sequencing collections

We sought *B. pseudomallei* whole genome sequences from the public database and combined these with newly sequenced genomes totalling 2,011, and further divided them into a discovery (1,136 isolate genomes) and a validation dataset (875 isolate genomes) (**Appendix 1 p5)**. Their accession numbers are tabulated in **Appendix 2**. The two datasets overlap geographically and temporally, spanning two major melioidosis endemic regions of Southeast Asia and Australia. These isolates came from environmental, animal and human sources with the latter constituting the larger proportion due to availability of microbiology laboratories embedded in the clinical settings. However, over 91% of human cases represent recent acquisition from the environmental sources thereby reducing the chance of co-evolutionary signals being shaped by human infection alone. For newly sequenced data, DNA libraries were sequenced on an Illumina Hiseq2000 with 100-cycle paired-end runs. Short reads were mapped to the reference *B. pseudomallei* genome K96243 with single nucleotide polymorphisms (SNPs) called as in.^6^ For the discovery dataset (n = 1,136), the alignment contained 389,476 SNPs, of which 206,019 and 183,457 were located in chromosome I and II, respectively. For the validation dataset (n = 875), the alignment contained 285,543 SNPs, of which 150,499 and 135,044 SNPs were located in chromosome I and II, respectively.

### Co-selection tests

Co-selection analysis was performed on the sequence alignment using the mutual information based GWES tool SpydrPick (**Appendix 3**).^7^ SNPs with minor allele frequency greater than 1% and gap frequency smaller than 15% were included in the analysis. To adjust for the population structure, sequence reweighting was applied using the similarity threshold of 0· 10. Direct links for which the mutual information exceeded the extreme outlier threshold, also after removing the influence of gaps, were selected for further examination. SNP-SNP pairs identified by the GWES were mapped by their coordinates to the K96243 reference genome. Most of the detected SNP-SNP pairs were located either within the same gene or in close physical proximity (**Appendix 1 p6**); an observation driven by linkage disequilibrium (LD). These pairs were separated by small physical distances and are unlikely to have had recombination events between them, resulting in so-called LD-mediated links. The LD decayed when the distance between the sites increased, and most SNPs were approximately in linkage equilibrium when each site was on a separate chromosome. While pairs located in close proximity are most likely explained by LD structure, some of them may represent genuine *cis* interacting partners. These may include SNPs in regulatory regions upstream of genes, such as DNA-binding sites for RNA polymerase and transcription factors; and non-coding RNA (ncRNA) which predominantly acts in *cis*. We thus grouped co-selected pairs into *cis* and *trans* based on the length of transcription fragments reported in *B. pseudomallei*.^8^ Here, pairs located on different genes or ncRNA but located within 7· 68 kb (a size covering 95^th^ percentile of polycistronic mRNAs) were termed *cis* pairs, while pairs located further than 7· 68 kb apart or on separate chromosomes were termed *trans* pairs. This resulted in *cis* or *trans* co-selection of gene-gene, gene-ncRNA, and ncRNA-ncRNA pairs. SNP-SNP pairs located within the same gene, or ncRNA were removed for this analysis.

### Expression patterns of genes and ncRNA under co-selection

We used *B. pseudomallei* condition-specific expression comprising 165 array profiles generated from Ooi *et al* to elucidate the function of co-selected gene-gene pairs (**Appendix 1 p5**).^8^ The data had been log-transformed to fit a Gaussian distribution. Expression correlations between gene-gene pairs were defined by Pearson correlation. We applied Benjamini-Hochberg adjusted correction for multiple testing. Expression profiles were categorised into 5 major categories spanning general growth, physical stresses, chemical stresses, infection and mutant conditions. For all conditions, and each of five major expression conditions; we tested if there was a significant difference in the correlated expression between co-selected gene-gene pairs and randomised gene-gene pairs using a nonparametric two-sided Mann-Whitney U test. Individual tests were performed for both discovery and validation datasets, and for both *cis-* and *trans* categories. Additional patterns of ncRNA expression under nutrient limitation were sought from Mohd-Padil *et al*.^9^ The authors measured and compared ncRNA expressed when *B. pseudomallei* was subjected to nutrient rich BHIB media and a nutrient-depleted M9 condition.

### Functional characterisation of a gene at the hub of co-selection

Construction of a gene knockout mutant for *BPSL1661*, a gene located at the co-selection hotspot, is described fully in **Appendix 1 p10**. We compared the growth- and stationary-phase survival of wild type (K96243) and *BPSL1661* mutant (K96243 Δ*BPSL1661*) across the conditions under which *BPSL1661* was reported to be upregulated (Ooi *et al*.), including nutrient limited media and high acidity. Details of the condition setup are described in **Appendix 1 p2-3**. For each condition, the bacterial population was enumerated at different time intervals. Differences in the growth profiles of wild type and mutant were determined using the Kolmogorov-Smirnov test.

## Results

### Co-evolutionary signals and their functional conservation

Our search for co-selection signals in the discovery dataset resulted in 13,061 SNP-SNP pairs spanning chromosome I (5,550 pairs), chromosome II (7,309 pairs), and between chromosome I and II (202 pairs); of which 8,035 pairs (61· 5%) could be replicated in the validation dataset. The congruence of the co-selection signals in the discovery and validation datasets is shown in **Appendix 3**, demonstrating that these complex patterns are common even across different datasets. The co-selected SNPs spanned genes, ncRNA and intergenic regions, each of which accounted for 69· 5%, 6· 8%, and 23· 7% of the total signals, respectively. This resulted in 334 *cis-* and 252 *trans* gene-gene pairs, 64 *cis-* and 11 *trans* gene-ncRNA pairs, and 19 *cis* ncRNA-ncRNA pairs from the discovery data; of which 383 pairs (56· 3%) were replicated in the validation data (**Figure 1, Appendix 4**). Despite *cis* pairs being more prevalent, the number of interacting partners per gene was fewer than the number of *trans* interacting partners (a mean of partners per *cis*-gene = 1· 11, a mean of partners per *trans*-gene = 2· 50) with the highest number observed in *BPSL1661* (*trans*-gene partners = 104 (discovery data), and 52 (validation data)).

**Figure 1.**
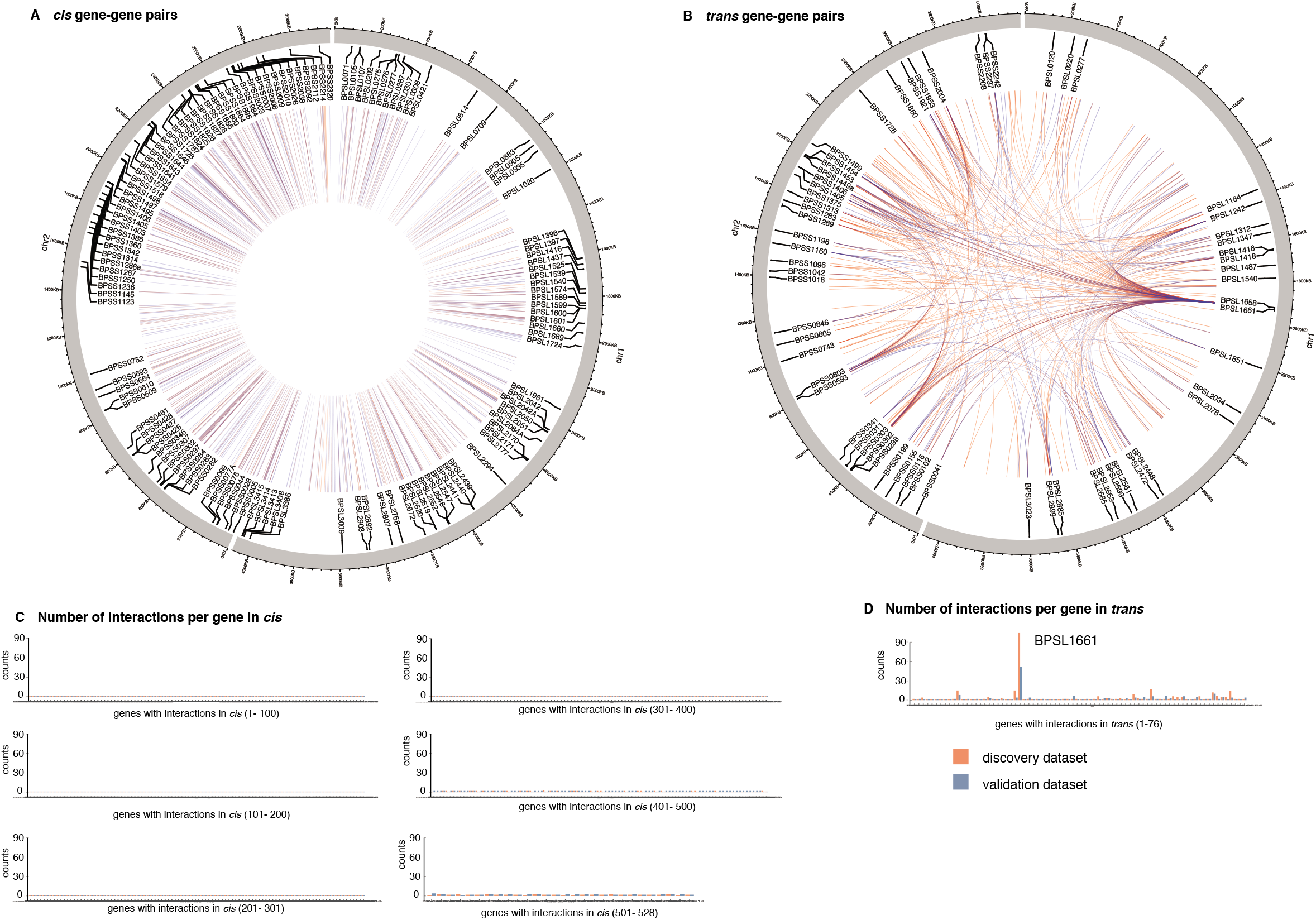
Co-selected gene-gene pairs in discovery and validation datasets. A, B) circos diagrams showing co-selected gene-gene pairs in *cis* (A) and *trans* (B). Any pairs located within 7.68 kb (95th percentile of transcription fragments) were categorised as *cis*, while those located further apart were grouped as *trans*. C, D) bar charts summarising the number of interacting partners per gene for *cis* (C) and *trans* (D) linkage. Data from discovery and validation datasets were highlighted in red and blue, respectively. A greater resolution figure can be accessed through https://figshare.com/s/82137a9c7427d8224d38

We assigned functional annotations to genes identified in both *cis* and *trans* gene-gene pairs. For both the discovery and validation dataset, the majority of the co-selected genes were functionally assigned as cell envelope (21· 9%), or transport/binding protein (10· 6%), while many were left as uncharacterised or hypothetical proteins (26· 7%) (**Appendix 1 p7**). We observed that a high proportion of the co-selected gene-gene pairs share the same functional annotations (51· 3% of *cis-* and 19· 9 % *trans* gene-gene pairs). To test whether this observation was driven by chance, we generated a separate randomised dataset for “*cis*” and “*trans*” gene-gene pairs using the same distance threshold (n = 1,000 pairs each) and compared the number of pairs under the same or different functional annotations between the randomised and real datasets. Excluding pairs with ambiguous annotations, functional conservation of *cis* gene-gene pairs followed random expectation (**Appendix 1 p7**). A possible explanation is that bacterial genes are organised into operons to control protein stoichiometry and maximise efficiency in protein production and function^10^, thereby increasing the chance of genes located in close physical proximity sharing the same functional annotation. However, conservation of functional annotation of *trans* gene-gene pairs was not random (**Appendix 1 p7**, two-sided Fisher exact test for discovery and validation datasets p-val < 1· 38 x 10^−2^). Our results indicate that co-selection of genes, at least in *trans*, is driven by gene function.

### Expression patterns of co-selected gene-gene pairs

We explored the expression profile of co-selected gene-gene pairs using *B. pseudomallei* whole genome tiling microarray expression data generated by ^8^. Expression profiles were assayed under a broad spectrum of conditions and exposure, including general growth (32 conditions), exposure to physicochemical stress (33 conditions), invasion assays (4 conditions), and defined genetic mutants (13 conditions) (**Appendix 1 p5**). Here, genes expressed in ≥ 70 conditions are defined as being constitutively expressed as in ^8^. Approximately 22· 4% of the genes that were detected in co-selection pairs in the discovery analysis or validation analysis, or both, were constitutively expressed compared to 39· 5% of genes that were not part of any co-selection pairs (two-tailed Fisher’s exact test p = 5· 74 x 10^−7^), indicating that co-selection signals were more strongly associated with condition-specific genes than those constitutively expressed.

We tested if *cis*- and *trans* gene-gene co-selection pairs also showed expression correlation compared to any randomised *cis*- and *trans* gene-gene pairs (n random pairs = 1,000 each). In addition, we examined under which conditions such correlations were observed. Using normalised gene expression profiles ^8^, Pearson’s correlation analysis was performed for each gene-gene pair with transcription data from all conditions tested as well as from subsets of those conditions. For all conditions and 4/5 tested condition subsets, co-selected genes paired in *cis* showed greater expression correlation than randomly paired *cis*-genes (**Figure 2A**, two-sided Mann-Whitney U test p values ≤ 1· 02 x10^−2^). The result suggests that expression of *cis* gene-gene pairs under co-selection are co-regulated. This is possibly mediated by the bacterial operon structure which enables the coupling of DNA transcription and messenger RNA translation, thereby enhancing the efficiency of protein production under each subset of conditions.^10^ An absence of operon structure for distally located genes should lead to less expression correlation for *trans* gene-gene pairs. Nevertheless, we observed greater expression correlation of co-selected genes paired in *trans*, than randomised controls, under physical stress conditions (**Figure 2B**, two-sided Mann Whitney U test p values ≤ 1· 05 x10^−2^), suggesting that physical stress has largely shaped the *tran*s co-selection patterns at the time-scale considered for this population. Here, physical stress conditions included temperature stress, osmotic stress, UV irradiation and nutrient deprivation; the environmental conditions *B. pseudomallei* is regularly or seasonally exposed to.^5,11–15^ However, it should be cautioned that *trans* gene-gene pairs are heavily driven by a few genes linked to many other genes. The expression profile or function of these few genes may thus largely drive the results observed here.

**Figure 2.**
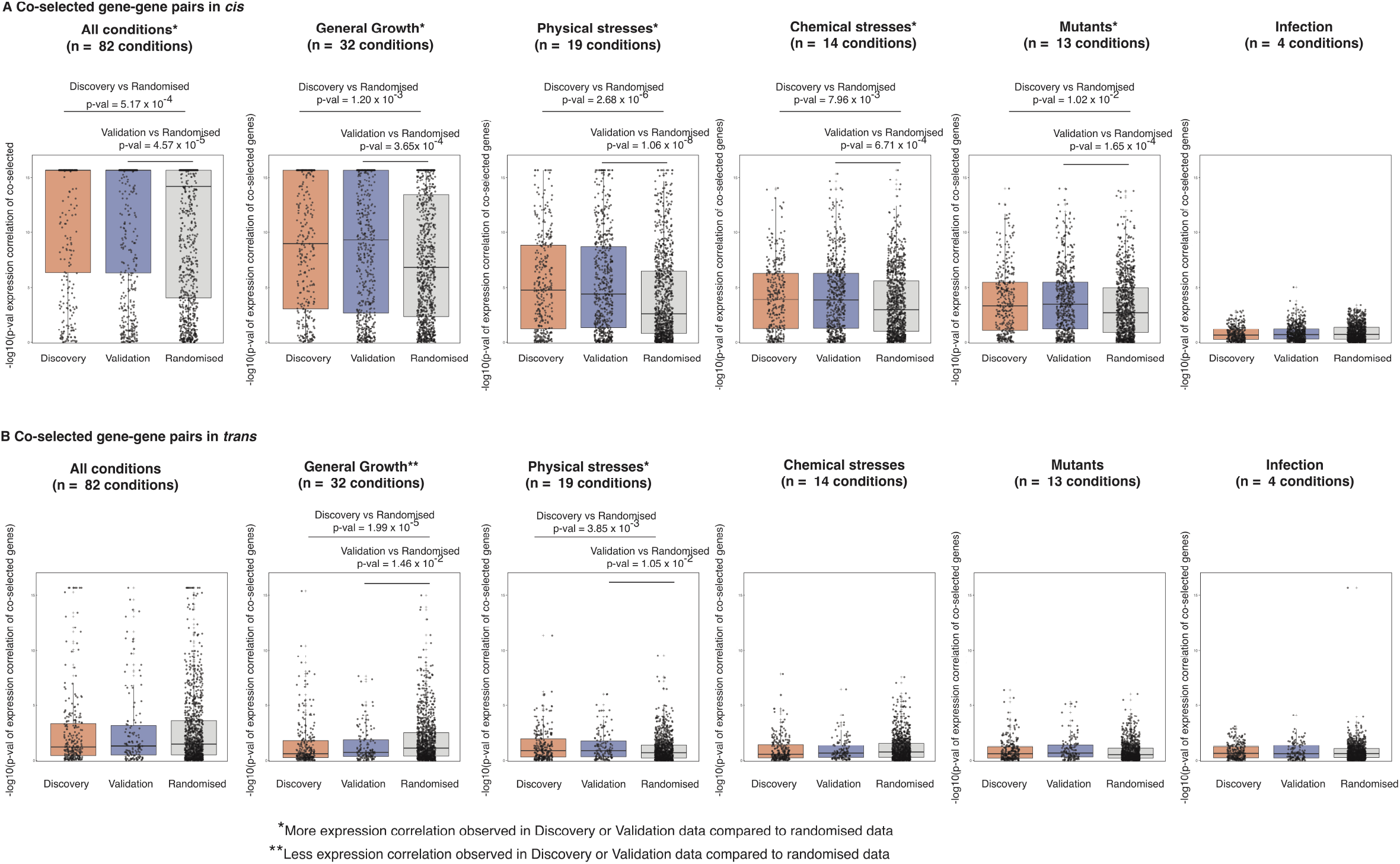
Expression correlation of co-selected gene-gene pairs. A) & B) represent expression correlations of co-selected gene-gene pairs from the *cis*- and *trans*- interactions, respectively. For all plots, vertical axis denotes the strength of expression correlation on negative log p-value scale while horizontal axis represents groups of data including true co-selection signals found in discovery (red), validation (blue) or randomised dataset (grey). The distribution of significant expression correlation of either discovery or validation data was compared to randomised data using two-sided Mann Whitney U test. The expression conditions tested (from left to right) include all conditions (n = 82), general growth (n = 32), physical stresses (n = 19), chemical stresses (n = 14), mutants (n = 13) and infection (n = 4). Each boxplot summarises first quartile, median, and third quartile with individual value plotted as a dot. A greater resolution figure can be accessed through https://figshare.com/s/0eed455b49b194a9840f

### A co-selection hotspot *BPSL1661* mediates bacterial survival under nutrient deprivation

Intriguingly, a putative adhesin gene *BPSL1661* constitutes 41· 3% and 38· 5% of *trans* gene-gene pairs in the discovery and validation datasets, respectively (**Figure 1B & 1D**). *BPSL1661* codes for a secreted protein with a size ranging from 2,594 to 3,230 amino acids (approximately 325 kDa) in different isolates (**Appendix 1 p8**). The gene is located in a large highly variable genomic region termed Genomic Island 8 (GI8)^16^, proposed to be acquired through horizontal gene transfer^17^ consistent with the presence of multiple *BPSL1661* alleles observed in our study. We detected six major alleles of *BPSL1661* (n ≥ five isolates) consistent with previous reports on multiple protein epitopes.^18,19^ All *BPSL1661* alleles share a conserved hemolysin-type calcium binding domain which is common in proteins secreted through a type I secretion system, and two copies of the VCBS domain (a repeat domain found in proteins encoded by *Vibrio, Colwellia, Bradyrhizobium* and *Shewanella*) known to be involved in cell adhesion. Variations in the presence of bacterial Ig and flagellin domains in *BPSL1661* were observed in the studied population. Interestingly, previous studies reported heterogeneity in human immune responses to polypeptides generated from different *BPSL1661* alleles, ranging from null to strong antibody responses.^18–20^ Such disparity in host recognition of different *BPSL1661* alleles potentially suggests that the protein may not principally function in host cell invasion but play other significant roles in *B. pseudomallei* survival. Transcription assays further revealed that *BPSL1661* is downregulated during *in vitro* and *in vivo* infection but upregulated in acidic conditions (pH 4· 0), mid-logarithmic phase in minimal media, and nutrient deprivation (**Figure 3A**); further suggesting *BPSL1661* is involved with adaptation to stress.

**Figure 3.**
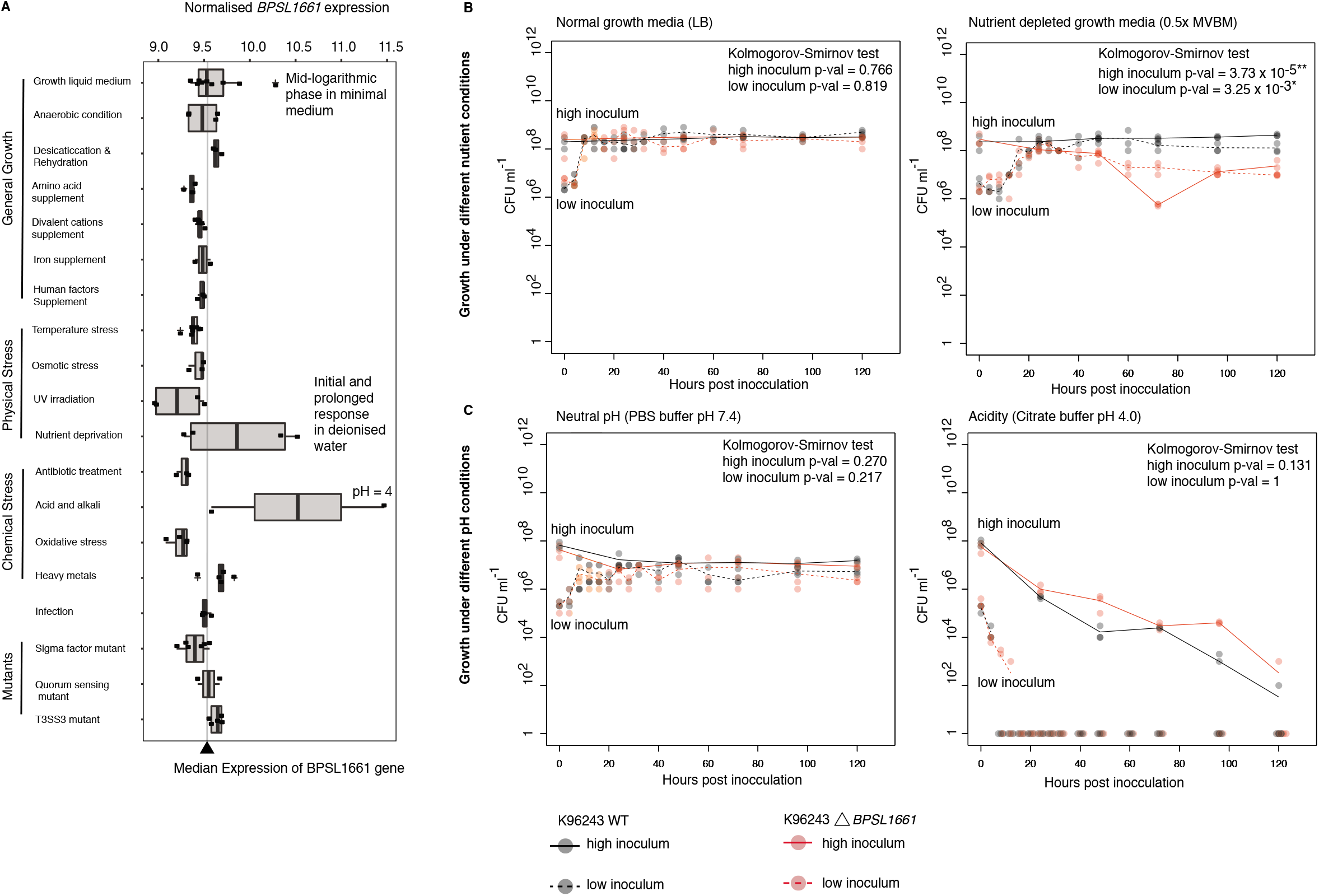
Functional characterisation of *BPSL1661*. A) Expression profile of *BPSL1661* across different conditions from Ooi *et al*. 2013, highlighting the gene upregulation during nutrient limited condition and high acidity. B) & C) represent growth- and stationary-phase survival of wild type and *BPSL1661* knockout mutant from K96243 reference strain under changes in nutrition (B) and pH (C). Bacterial survival post inoculation at low (10^6^ CFU ml^-1^) and high (10^8^ CFU ml^-1^) at different time intervals. Three replicates were taken at each timepoint. Black and red dots denote observations from wild type and *BPSL1661* mutant, respectively. Dotted and solid line represent profile of low- and high bacterial inoculum, respectively. Difference in growth profiles between wild type and mutant was measured using two-sided Kolmogorov-Smirnov test. A greater resolution figure can be accessed through https://figshare.com/s/f40dca8e693331773ebb

To better understand the function of *BPSL1661*, we knocked out the *BPSL1661* gene in the K96243 strain and compared the number of live cells of the mutant against the wild type, under the conditions in which *BPSL1661* was upregulated, for 120 hours (**Figure 3A**). We observed no significant differences in bacterial survival in nutrient rich growth media, neutral pH (pH 7· 4) or acidic conditions (pH 4· 0). This could be due to functional redundancy that compensates for the loss of a single gene function. However, the *BPSL1661* mutant showed a significant reduction in stationary-phase survival compared with the wild type under nutrient limited conditions (two-sided Kolmogorov-Smirnov test p-value = 3· 73 x 10^−5^ and 3· 25 x 10^−3^ for high and low bacterial inoculum, respectively); confirming an essential role of *BPSL1661* under nutrient deprivation. Our observation of *BPSL1661* as a hotspot may imply that nutrient depletion is the selective pressure underlying the co-selection patterns. Indeed, environmental sampling studies in Southeast Asia and Australia have consistently reported that the bacterium is commonly found in nutrient depleted soils ^5,13^. Although lower nutrient abundance appears to be a common feature across melioidosis endemic areas; the soil physiochemical properties, microbial diversity, temporal disturbances such as monsoon seasons and anthropogenic activities that alter the environmental conditions vary greatly between each area and between studies ^14,21–24^. These factors create patterns of spatial and temporal heterogeneity to which *B. pseudomallei* has adapted and possibly has led to the coexistence of multiple *BPSL1661* alleles detected in this study. We also noted geographical differences in *BPSL1661* allele frequencies. An allele harbouring a flagellin domain (here denoted as allele A, **Appendix 1 p8-9**) was detected at lower frequency in Australia (28· 7%), at moderate frequency in the Malay Peninsula (37· 6% from Malaysia & Singapore) and higher frequency in the countries bordered by the great Mekong river (59· 1 % from Thailand, Laos, Cambodia and Vietnam) (**Appendix 1 p8**). Such a difference in allele frequencies could be either driven by different local selection pressures, or by genetic drift (or both). Functional characterisation of different *BPSL1661* alleles also warrants further future studies.

### *BPSL1661* co-selection network and putative bacterial response under low nutrient abundance

We considered genes and ncRNA co-selected with *BPSL1661* in both the discovery (n gene pairs = 105, n ncRNA pairs = 5) and validation dataset (n gene pairs = 53, n ncRNA pairs = 2) totalling 136 pairs, of which 29 pairs are shared in both datasets (**Figure 4, Appendix 5**). The majority of these gene and ncRNA pairs were linked to *BPSL1661* in *trans* except for an outer membrane protein *BPSL1660* which paired in *cis*. During nutrient depleted conditions ^8,9^, 43/129 of the *trans* gene pairs and 1/6 of ncRNA pairs were upregulated with *BPSL1661*, while 64/129 of the *trans* gene pairs and 2/6 of ncRNA pairs were downregulated, respectively. Many of these genes are predicted to encode proteins that participate in alternative metabolic pathways, energy conservation, uptake of external carbon source, cellular signalling, and transcriptional regulation (**Appendix 5**). A pyrophosphohydrolase gene (*spoT* or *BPSL2561*) is upregulated during nutrient starvation and is known to have a dual function to synthesise and hydrolyse guanosine tetra- and pentaphosphates (ppGpp).^25^ In bacteria, ppGpp serves as a mediator in nutritional surveillance, coordinating a variety of cellular activities in response to changes in nutrient availability. ^26^ In *B. pseudomallei*, deletion of ppGpp synthetase and hydrolase led to reduced survival during stationary phase compared to wild type. ^25^ It is possible that *B. pseudomallei* switches to alternative carbon sources to maintain cellular energy when complex sources such as glucose are not available. Genes encoding a C4-carboxylate transport transcription regulation protein (*BPSL0427*), a malate synthase (*BPSL2192*) and a glycogen branching enzyme (*BPSL2076*) were co-selected with *BPSL1661* but had different expression profiles under nutrient limited conditions. A transport transcription regulation protein (*BPSL0427*) was upregulated under nutrient deprivation. Its homolog was shown to facilitate the cellular uptake of four carbon compounds such as aspartate, fumarate, and succinate when common carbon sources such as glucose are scarce.^27^ Another upregulated gene, malate synthase (*BPSL2192*) is a key enzyme involved in the glyoxylate cycle. Its expression allows cells to utilise two-carbon compounds to sustain carbon requirement in the absence of glucose.^28^ In contrast, a glycogen branching enzyme (*BPSL2076*) was co-selected with *BPSL1661* but downregulated during nutrient depletion. A homolog of the glycogen branching enzyme is known to facilitate glucose conversion into long term glycogen storage when there are excessive carbon sources.^29^ The co-selection of *spoT, BPSL0427, BPSL2192* and *BPSL2076* with *BPSL1661* may reflect the energy balance of cells under ranges of nutritional conditions through their evolutionary timeline. Together, the *BPSL1661* co-selection network seems to suggest that *B. pseudomallei* has adapted to survive nutrient limited conditions and/or hostile environments.

**Figure 4.**
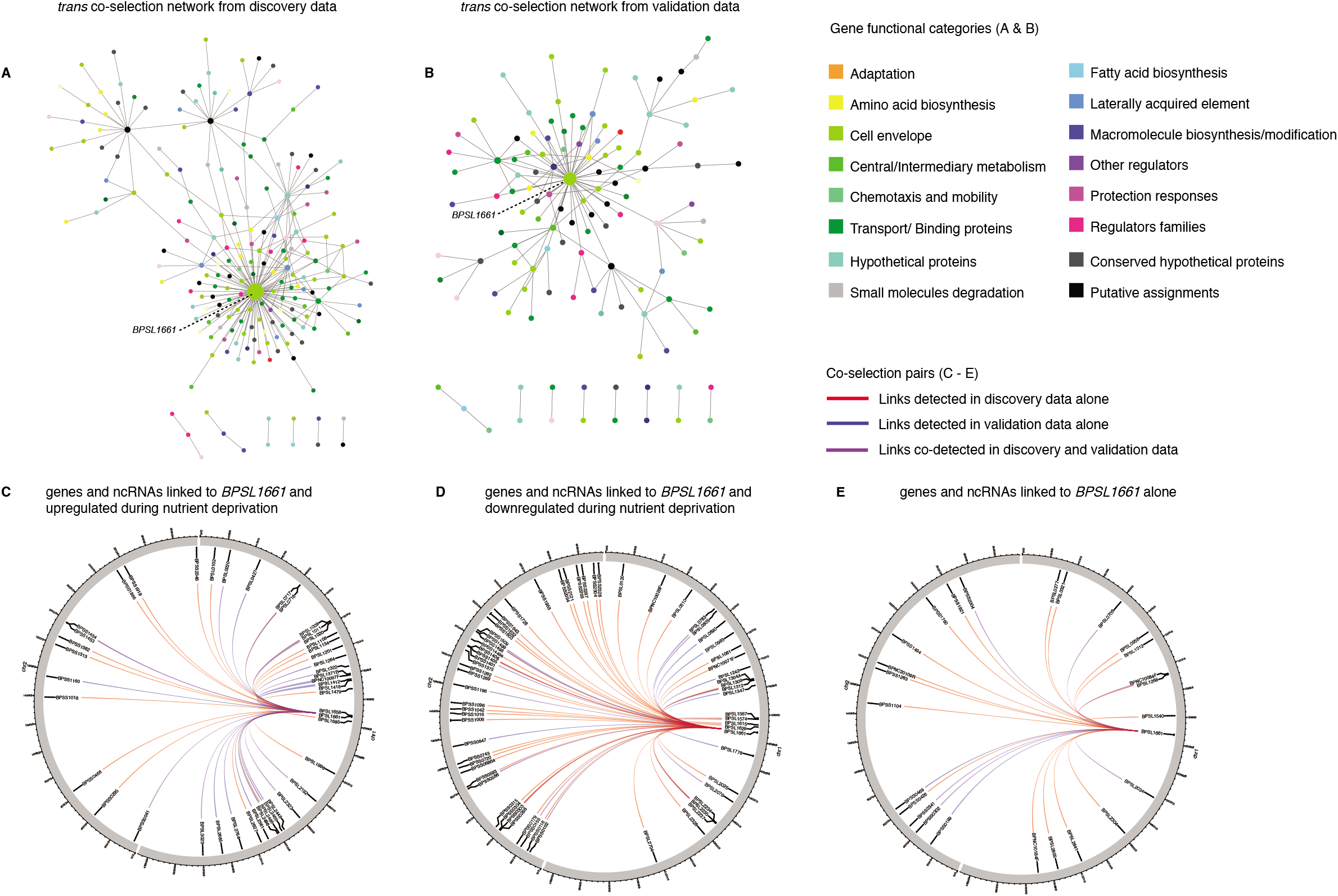
*BPSL1661* co-selected gene and ncRNA network. A) & B) represent networks of *trans* interacting co-selected gene-gene pairs in the discovery and validation datasets, respectively. Each node denotes a gene under co-selection with the node size proportional to numbers of pairs linked to the genes, and colour coded by the gene functional category. For both discovery and validation datasets, *BPSL1661* consistently acts as hub of the *trans* co-selected gene network. C), D) & E) summarise genes and ncRNA co-selected with *BPSL1661* in both discovery and validation dataset: genes and ncRNA co-selected and upregulated under nutrient deprivation (C); genes and ncRNA co-selected but downregulated under nutrient deprivation (D); and genes and ncRNA that are co-selected alone (E). Links identified from the discovery data alone, validation data alone, and both data are coloured as red, blue and purple, respectively. A greater resolution figure can be accessed through https://figshare.com/s/394c5e95abd58bdff091

## Discussion

This study is the first, to our knowledge, to deploy an integrated approach of GWES, transcriptomic analyses and knockout assays to understand the evolution and unique selective pressures acting on a micro-organism. While GWES detected signals for co-selected loci, transcriptomic data provided condition-dependent information on which selection pressures may have acted upon the detected loci. However, our study focused only on nucleotide polymorphisms found by comparison to the *B. pseudomallei* K96243 reference genome. Co-selected loci on other types of structural variants including indels, genomic inversions, gene duplication or horizontally acquired genes absent in K96342 will be missed from this analysis. It is also possible that other transcriptional conditions the bacterium is exposed to in its native niche are missing from our study. The incomplete genomic and transcriptomic data warrant further studies to cover more complex genomic variants, and broader transcriptional conditions. Nevertheless, our integration of data offers stringent predictions on which genes are key to *B. pseudomallei* survival under specific conditions. In particular, the putative adhesin *BPSL1661* was identified as a hotspot of the co-selection map. Our gene knockout experiment confirmed that the gene is essential for survival under nutrient deprivation. This is consistent with the soil conditions in which *B. pseudomallei* are commonly found and provides evidence for the hypothesis that unlike most bacteria, *B. pseudomallei* has a competitive advantage in nutrient depleted environments.

The presence of *B. pseudomallei* in nutrient-depleted soil defines geographical regions where humans are at risk of melioidosis. Several factors, including anthropogenic activities could lead to soil erosion and nutrient depletion. With greater land-use pressure, poor agricultural practice such as the shortening of fallow periods and more frequent cropping rotations as well as conversion of marginal land to cash crops increasingly occur throughout Southeast Asia and could lead to increased loss of soil nutrient. In melioidosis endemic area such as Australia, bushfires and an ancient firestick farming practiced by Indigenous people over the past 100,000-120,000 years have largely shaped the continent’s landscape and led to the spread of fire adapted species^32^. In South- and Southeast Asia which are another melioidosis endemic areas, crop residue burning after harvest is an ongoing practice and reported to link with reduced soil nutrients ^30,31^. Lower crop yields as a result of intense farming mean farmers need to clear further field and repeat similar cycles thereby expanding the nutrient-depleted areas. Farmers who could not afford more lands may switch to continuous cultivation of cash crops such as maize or cassava, which lower the soil nutrient and exacerbate the problem. Together, these practices could improve conditions for *B. pseudomallei* survival. It is therefore notable that we observe genes required for survival of nutrient depletion at the hub of the co-selection signals. Our study predicts soil nutrient depletion as improved conditions for *B. pseudomallei* survival and a corresponding increase in the incidence of melioidosis, and calls for improving environmental health to assist melioidosis prevention.

## Contributors

J.C. conceived the original concept. C.C., R.W.S. and J.C. designed the experiment and oversaw the project and analyses. C.C., S.J.P., N.P.J.D., J. Parkhill, N.R.T., R.W.S. and J.C. acquired funding. P.B., K.S., E.J.F., S.D., N.C., D.L., and N.P.J.D. contributed samples and reagents or assisted in samples preparation and permitting. J. Pensar and M.P. developed analytical tools. C.C. and J. Pensar performed genomic analyses. S.C. developed gene knockout mutant. S.C. and C.P. performed experimental validations. C.C., J. Pensar, A.S., E.J.F., N.C., D.L., J. Parkhill, N.R.T., R.W.S. and J.C. contributed to the interpretation and presentation of results in the main manuscript and supplementary documents. C.C. and J.C. wrote the first draft with input from all authors.

## Acknowledgement

The authors thank Dr Alain Pierret, Dr Olivier Ribolzi and Dr David Dance for insightful discussion on the environmental implication of the study. C.C. was funded by Wellcome International Intermediate Fellowship (216457/Z/19/Z) and Sanger International Fellowship. J.C. was funded by the ERC grant no. 742158. S.C., C.P. and R.W.S were funded by Targeted Research Grant, Faculty of Medicine, Khon Kaen University. This publication presents independent research supported by the Health Innovation Challenge Fund (WT098600, HICF-T5-342), a parallel funding partnership between the Department of Health and Wellcome Trust. The views expressed in this publication are those of author(s) and not necessarily those of the Department of Health or Wellcome Trust. This project was also funded by a grant awarded to Wellcome Trust Sanger Institute (098051), and to the Wellcome Thailand and African Programme (106698).

## Appendix1

### Supplementary Methods

#### Functional classification of genes under co-selection

SNP-SNP pairs identified in GWES were mapped by their coordinates to genes and ncRNA annotated in K96243 reference genome^1^, resulting in co-selection of gene-gene, gene-ncRNA, and ncRNA-ncRNA pairs. SNP-SNP pairs located within the same gene, or ncRNA were removed for this analysis. Different gene functional categories including a curated Riley’s classification^2^, clusters of orthologous groups (COGs)^2^, KEGG Pathway^3^, and gene ontology (GO terms)^4^ were assigned to each gene-gene pair. The final analysis was focused on a curated Riley’s classification as it covered more genes detected for co-selection than other classification systems. For both discovery and validation datasets, we searched for enrichment of gene functions among co-selected genes against their distribution in K96243 genome using two-sided Fisher’s exact test (R function fisher.test()) while controlling for false positive from multiple testing using Benjamini-Hochberg method^5^.

We next tested for functional dependency of co-selected gene-gene pairs by comparing the frequency of pairs with the same functional annotations to pairs with different functions. To test whether this was observed by chance, we simulated a set of 1,000 random gene-gene pairs from the K96243 genome with co-selected genes excluded. A separate randomised collection was pooled for *cis* (genes located < 7.68 kb apart) and *trans* interactions (genes located > 7.68 kb apart). Two-sided Fisher’s exact test was separately performed to compare the distribution of co-selected pairs with the same and different gene functions against randomised datasets from both discovery and validation datasets. Given that the bacterial genes are organised into operons, the test was also separately conducted for *cis-* and *trans* interactions where paired genes are likely under the same and different operons, respectively.

#### Genetic variations in *BPSL1661* and their distribution across the core genome phylogeny

Where complete genomes were available, BLAT v. 36^10^ was used to locate the position of *BPSL1661* homolog and further confirmed with genome annotations. Illumina sequenced short reads were assembled as in^11^ and annotated using Prokka v.1.14.5.^12^ Coding sequence of *BPSL1661* was identified using BLAT and confirmed with gene annotation. *BPSL1661* sequences from all genomes were aligned using MAFFT v.7.407 ^13^ and assigned into different clusters using CD-HIT-EST v.4.8.1^14^ with sequence identity threshold of 0.9. Here, any clusters with ≥ 5 members were considered representative alleles, resulted in 6 *BPSL1661* alleles across the datasets. Protein domains of each *BPSL1661* allele were sought from CDD/SPARCLE v.3.17 conserved domain database^15^.

To investigate the distribution of *BPSL1661* variants across *B. pseudomallei* population, core-genome phylogeny was constructed from core genome SNP alignment. SNPs were called from K96243 mapped genome alignment using SNP-sites v.2.5.1^16^ excluding sites associated with mobile genetic elements.^17^ We next estimated a maximum likelihood tree using IQ-TREE v.1.6.10 ^82^ using General Time Reversible (GTR) + Gamma distribution model of nucleotide substitution with default heuristic search options and 1,000 bootstraps^18^.

#### *BPSL1661* functional characterisation

The hub of co-selection signals, *BPSL1661* was next functionally characterised. Bacterial strains, plasmids and oligonucleotides used in this study are listed in **Supplementary Table 1**. GF-1 bacterial gDNA extraction kit and deoxynucleotide triphosphates (dNTPs) were purchased from Vivantis; Platinum DNA Taq polymerase from Invitrogen; pGEM-T Easy vector systems from Promega; KOD Plus DNA polymerase from Toyobo; Restriction enzymes from New England BioLabs; QIAquick Gel Extraction kit, MinElute PCR purification kit from Qiagen; Ampicillin (Ap), Kanamycin (Km), Gentamycin (Gm) were purchased from Sigma, and Isopropyl β-D-1-thiogalactopyranoside (IPTG), 5-bromo-4-chloro-3-indolyl-β-D-glactopyranoside (X-Gal) and 5-bromo-4-chloro-3-indolyl-β-D-glucuronide (X-Gluc) were purchased from Gold Biotechnology.

The culture of *B. pseudomallei* K96243 wild-type, *B. pseudomallei BPSL1661* clean mutant, and the *Escherichia coli* strains used for construction of the *B. pseudomallei* mutant were routinely grown in Luria– Bertani (LB) and Luria-Lennox (LB, low-salt) medium (Sigma, USA), at 37°C with 200 rpm agitation. When necessary, the medium was supplemented with antibiotics, chemicals and chromogens at the concentrations of 100 µg ml^-1^ Ap, 35 µg ml^-1^ Km, 0.1 mM IPTG, 50 µg ml^-1^ X-Gal for *E. coli*; 5 µg ml^-1^ Gm, 1,000 µg ml^-1^ Km, 0.1 mM IPTG, 50 µg ml^-1^ X-Gluc for *B. pseudomallei*.

#### Construction of the *BPSL1661* clean deletion mutant

The nucleotide sequence encoding (*BPS_RS08795*, [*BPSL1661*]) gene (GenBank accession no. WP_045606470.1) of *B. pseudomallei* K96243 was used to design primers for clean deletion using Primer-BLAST program. The BPSL1661 clean mutant (Δ*BPSL1661*) (**Supplementary Figure 6**) was constructed from *B. pseudomallei* K96243 by double-crossover allelic exchange as described previously. Briefly, the upstream and downstream DNA fragments of *BPSL1661* were amplified using BPSL1661_PFup/PRup and BPSL1661_PFdown /PRdown primers and then ligated (**Supplementary Table 1**). The ligated amplicons were cloned into TA cloning vector pGEM-T Easy (Promega, USA) and check for its correct insert size before subcloned into an allelic exchange plasmid pEXKM5.^19^ The recombinant pEXKM5 plasmids were introduced into *B. pseudomallei* K96243 by bi-parental conjugation using conjugal *E. coli* S17-1pir.^20^ Merodiploid was selected on LB medium supplemented with X-Gluc and Kanamycin that appeared as pale blue colonies. To obtain the clean deletion Δ*BPSL1661* mutant, merodiploid colonies were cultured in LB medium to reach stationary phase then subculture into yeast extract tryptone (YT) broth supplemented with 10 % sucrose for overnight and spread on LB-medium with 15% sucrose. Suspected *B. pseudomallei* colonies was verified by PCR and confirmed by DNA sequencing of the region flanking *BPSL1661*.

#### *BPSL1661* condition-specific assays

Guided by expression profile of *BPSL1661* in Ooi *et al*^2^, the growth of wild-type and Δ*BPSL1661* were compared by enumerating the bacterium grown under nutrient rich medium (LB broth), nutrient depleted condition (MVBM), neutral pH and acidic pH. The bacterium was grown on Ashdown’s selective agar^21^ before transferring to grow under each condition.

We followed Ooi *et al*^2^ for normal growth condition using LB broth. We artificially induced nutrient limited condition *in vitro* by replacing glucose limited media (six carbon source) with glycerol (three carbon source) in MVBM (Modified Vogel and Bonner’s medium). Morever, MVBM^22^ further forced cell starvation by inducing biofilm formation. Nutrients will be consumed by cells located on the periphery of the biofilm clusters leading to reduced level of nutrients diffused to the inner cells. Both wild-type and Δ*BPSL1661* were grown on 0.05% glucose in 0.5 x MVBM [0.05%G, 0.5×MVBM]) at 37°C with 200 rpm shaking for overnight. The culture was adjusted and measured an optical density (OD) at OD600 nm to be 0.2 (OD600 = 0.2) and then inoculated into 10 ml prewarmed 0.05% glycerol, 0.5×MVBM medium at 37°C with 100 rpm shaking.

Growth in neutral pH was conducted in 10-fold diluted phosphate buffered saline (PBS) pH 7.4. Growth in a low pH was done as described previously^2^. For both condition, bacteria were cultured for overnight in LB broth at 37°C with 200 rpm shaking and sub-cultured in new fresh medium until log phase. Bacterial cells were washed in 1×PBS buffer pH 7.4 or 1×citrate buffer pH 4.0 for three times by centrifugation (8000 × g for 5 min) and resuspended in 10 ml of each buffer.

For all conditions, we observed bacterial initial growth and stationary-phase survival for both low (10^6^ CFU ml^-1^) and high (10^8^ CFU ml^-1^) inoculum. The colony forming unit (CFU) post inoculation was counted at different time interval. All assays involved three replicates that were independently prepared, cultured and treated. Difference in growth profile between wild-type and Δ*BPSL1661* for each condition was compared by a nonparametric Kolmogorov-Smirnov test.

**Supplementary table 1.**
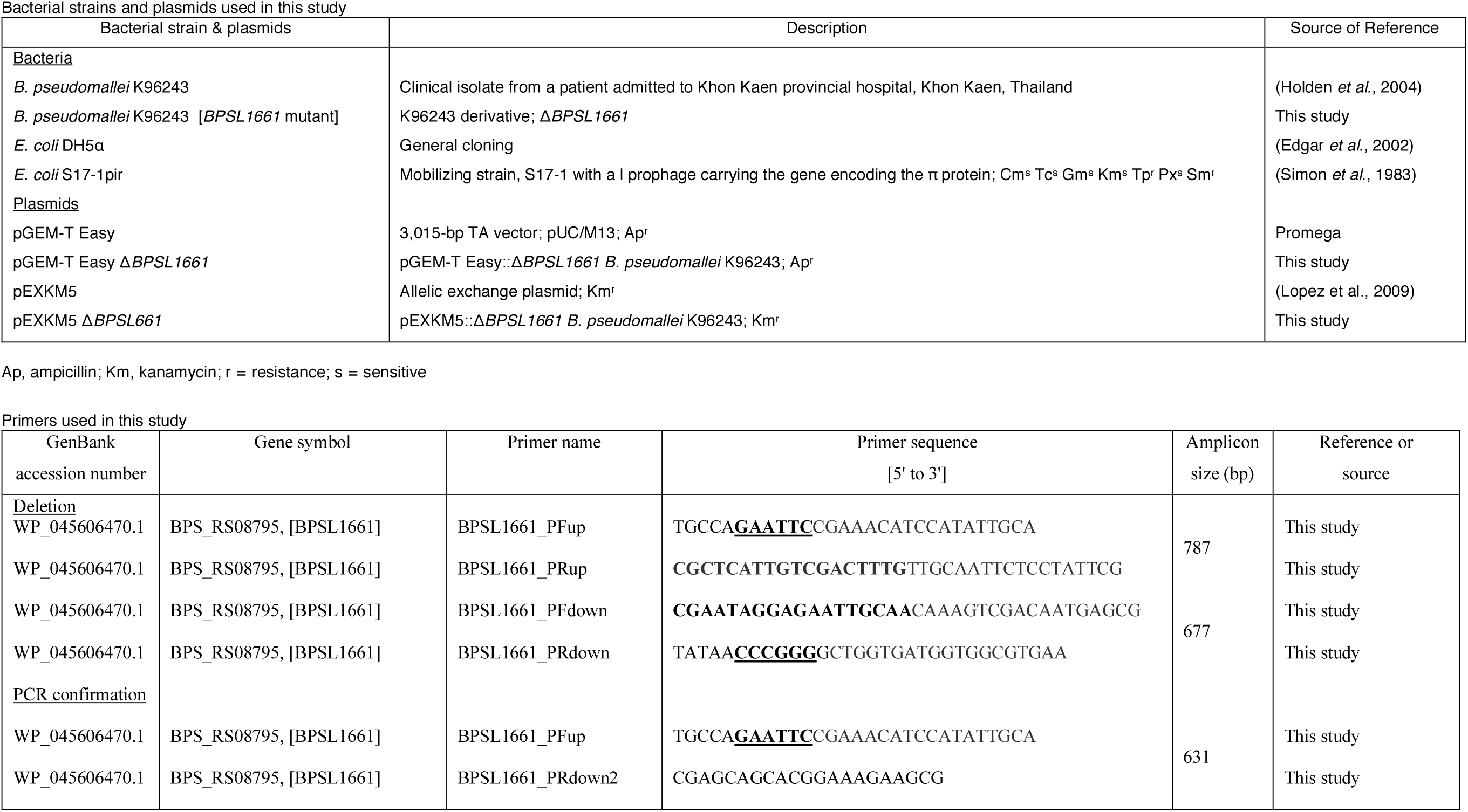

**Supplementary Figure 1.**
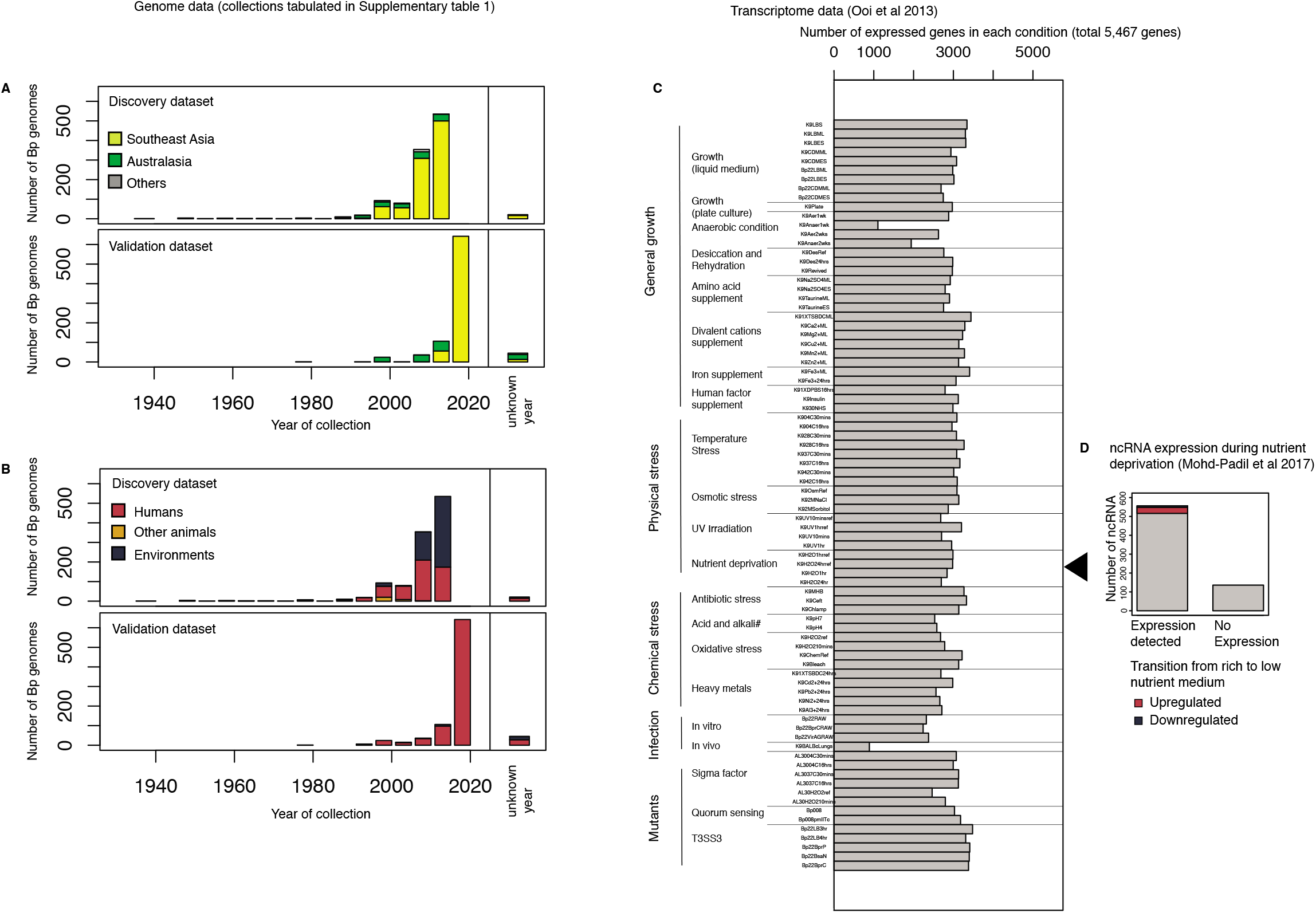
Summary of genomic and transcriptomic data used for the analyses. A) - B) represent distribution of *B. pseudomallei* genomes in the discovery and validation dataset by year of collection with geographical origins and source of isolates plotted in A) and B) respectively. c) summarises *B. pseudomallei* expression profile across 82 conditions covering general growth, physical stress, chemical stress, infection and mutants obtained from Ooi *et al*.^2^ The bar chart highlights the number of genes expressed in each condition. D) represents the expression profile of *B. pseudomallei* non-coding RNA in nutrient-rich and nutrient-limited conditions. The data was obtained from Modh-Padil *et al*.^9^ (A high resolution figure is also archived in https://figshare.com/s/3ee6203d74d5b37109ab)

**Supplementary Figure 2.**
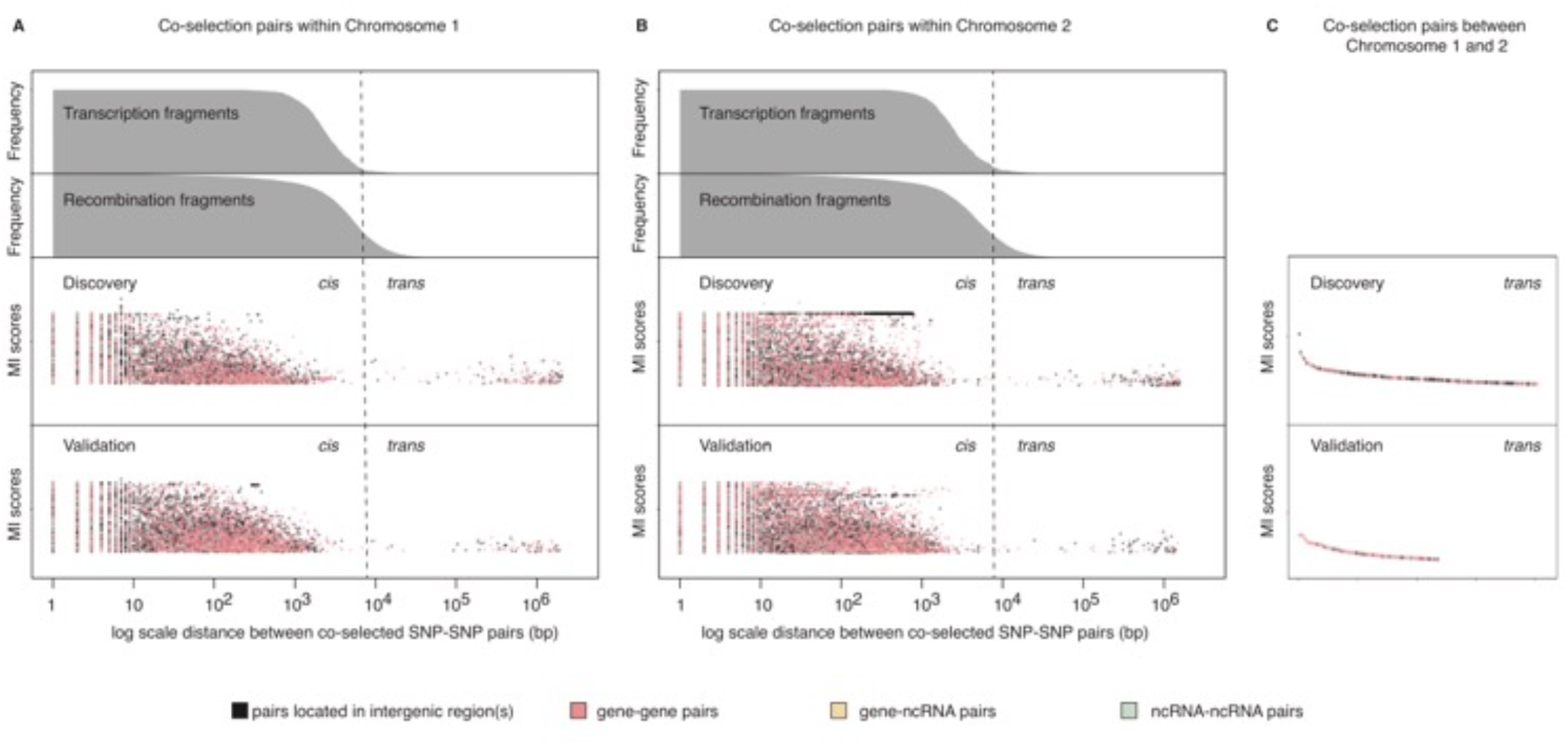
Co-selected SNP-SNP pairs. A) - C) represent co-selection sites detected on Chromosome 1, Chromosome 2, and between Chromosome 1 and 2, respectively. For A) & B) The horizontal axis denotes physical distance on the chromosome on the log10 scale. The top to bottom panels represent the accumulated size of transcription fragments, the accumulated size of recombination fragments, the distance between co-selected SNP-SNP pairs against the mutual information scores from the discovery datasets, and the validation datasets, respectively. For two bottom panels, each dot represents a co-selected SNP-SNP pair and is colour-coded by their associations with genes (pink), a gene and a molecule of non-coding RNA (orange), molecules of non-coding RNA (green), or intergenic regions (black). Vertical dotted lines denote 95^th^ percentile of transcription fragments. C) Top to bottom panel represent the mutual information scores of each co-selected SNP-SNP pair identified in the discovery and validation datasets, respectively. The same colour scheme was employed as in A) & B). (A high resolution figure is also archived in https://figshare.com/s/ef63fa3697532fab3f26)

**Supplementary Figure 3.**
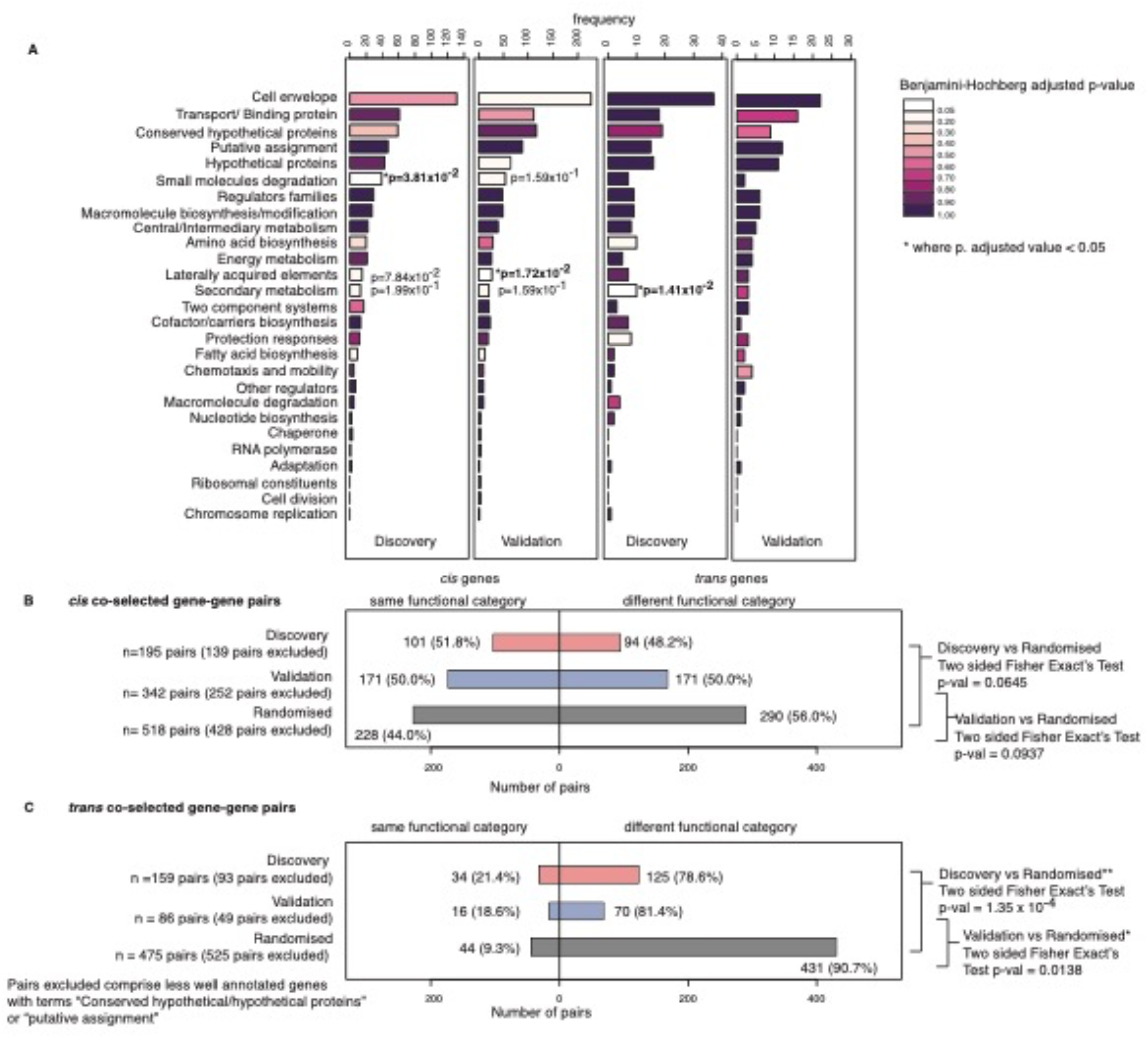
Functional category of co-selected genes. (A) Enrichment of functional category observed for *cis*- and *trans*-interactions in both discovery and validation datasets. The colour indicates the significance of the enrichment test result, ranging from white (adjusted p-value <0.05) to dark purple (adjusted p-value = 1). B) & c) represent distribution of co-selected gene-gene pairs with same (left) and different (right) functional annotations observed for *cis*- (B) and *trans-* (C) interactions. Data obtained from the discovery, validation, randomised dataset were highlighted as red, blue, and grey, respectively. The distribution of pairs with same and different functional annotations were compared against the randomised dataset using two-sided Fisher’s Exact Test. Pairs with ambiguous annotations comprising terms “conserved hypothetical/hypothetical proteins” or “putative assignment” were excluded from the analyses. (A high resolution figure is also archived in https://figshare.com/s/63c45bebcfa0a42800a6)

**Supplementary Figure 4.**
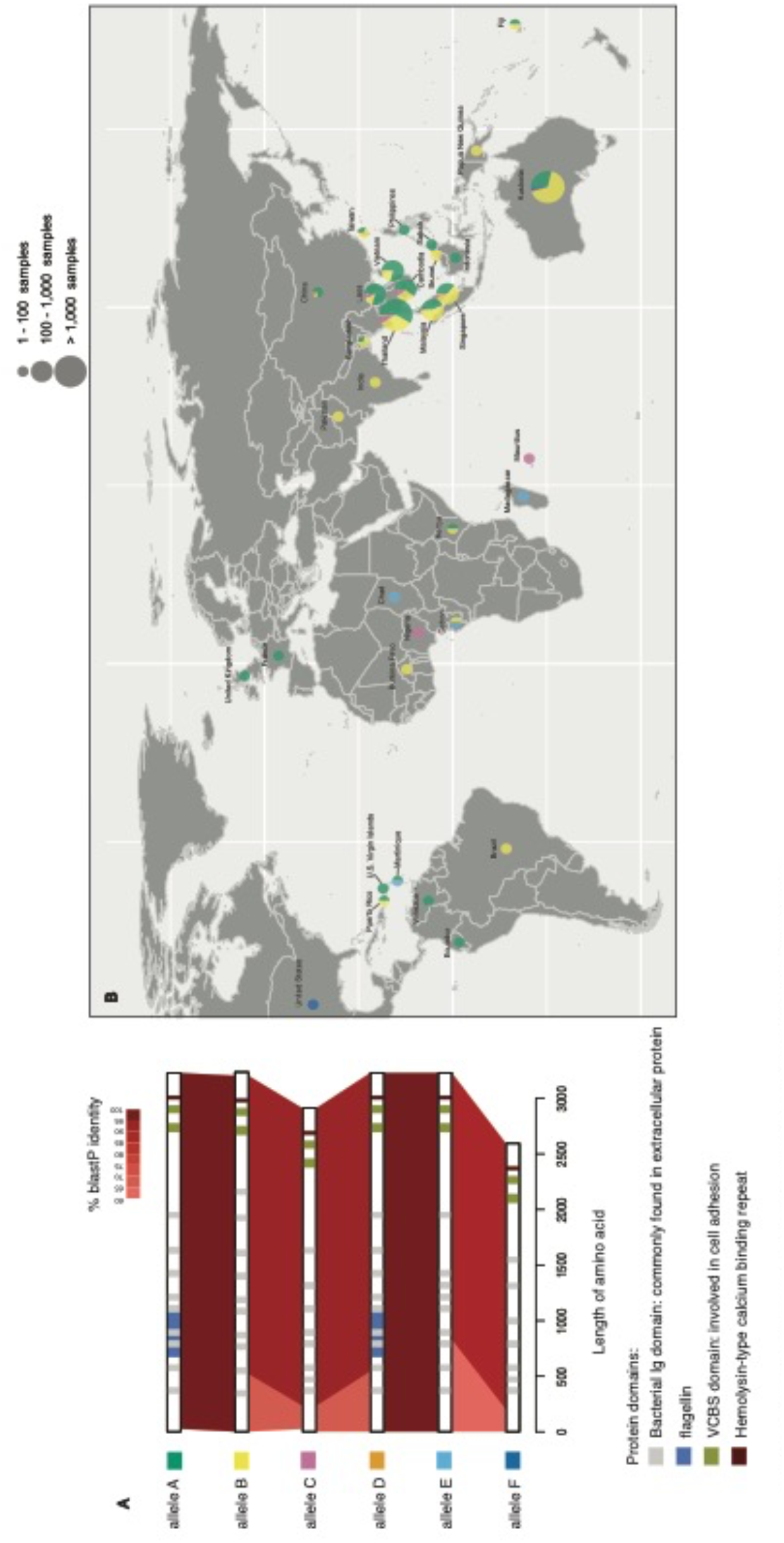
Distinct *BPSL1661* alleles and their geographical distribution. A) represents diagram of six different *BPSL1661* alleles (denoted A to F). Each allele harbours different combination of protein domains including bacterial Immunoglobulin (Ig) domain (grey); flagellin (blue); *Vibrio, Colwellia, Bradyrhizobium*, and *Shewanella* domain (VCBS, green); and hemolysin-type calcium binding repeat (brown). Red areas show protein identity. B) a world map shows geographical distribution of different *BPSL1661* alleles. Colour in pie charts correspond to different BPSL1661 alleles, with the size of the pie charts proportional to number of samples obtained from each area. (A high resolution figure is also archived in https://figshare.com/s/cf8d8357235744c4050f)

**Supplementary Figure 5.**
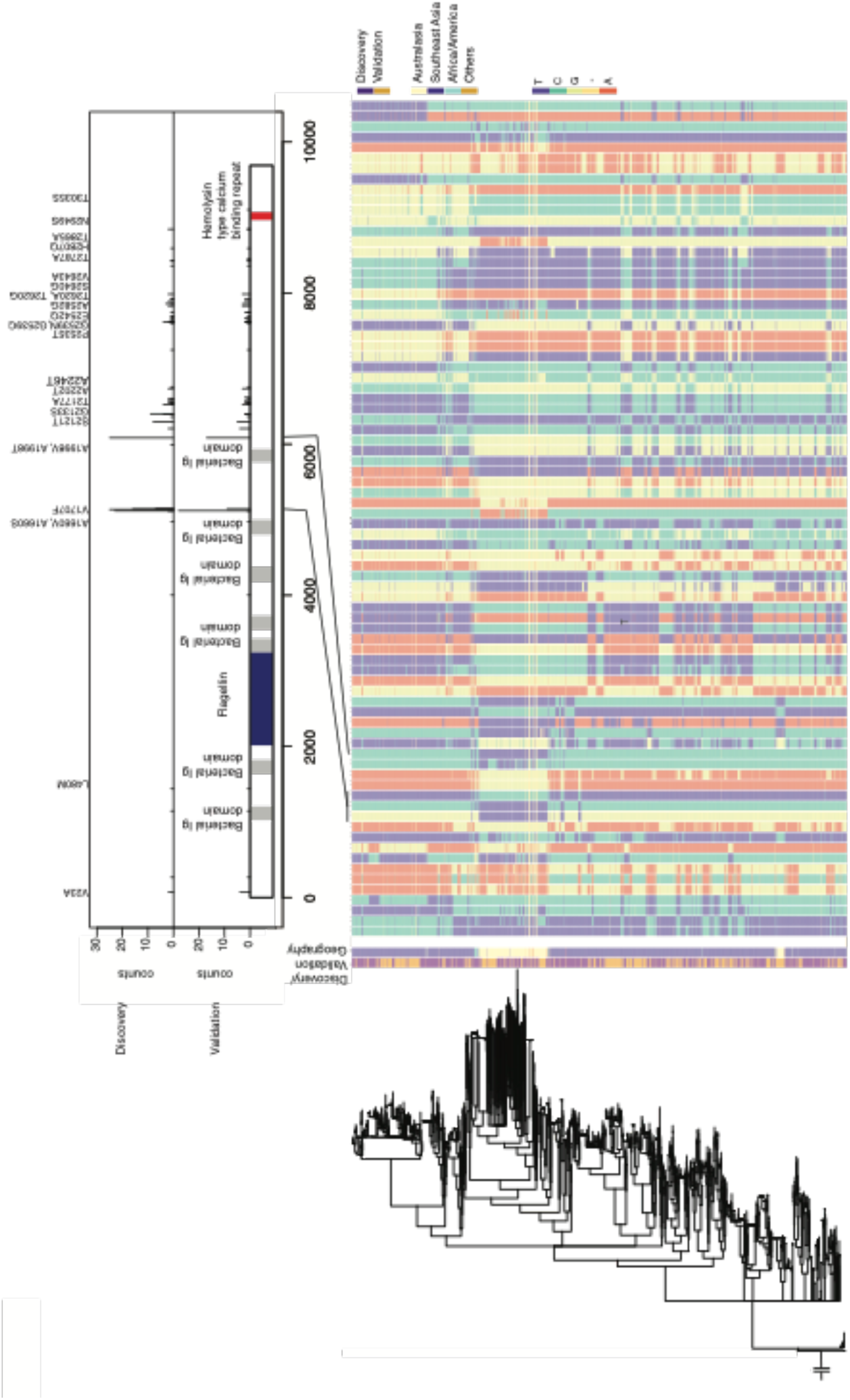
Distribution of *BPSL1661* variants. a) highlights SNP locations on *BPSL1661* where coevolutionary signals were detected in discovery (top) and validation datasets (bottom). Peaks corresponding to nonsynonymous substitutions were labelled. b) represents distribution of alleles by *B. pseudomallei* core-genome phylogeny. The estimated phylogeny is shown on the left with the first two columns labelled by type of data (discovery or validation dataset) and geographical origins of isolates, respectively. The remaining columns demonstrate all nucleotide variants detected in *BPSL1661*. Variants coincide with peaks of coevolutionary signals were marked, one of which correspond to V1707F substitution. (A high resolution figure is also archived in https://figshare.com/s/92b0485f1c7cb4658bc9)

**Supplementary Figure 6.**
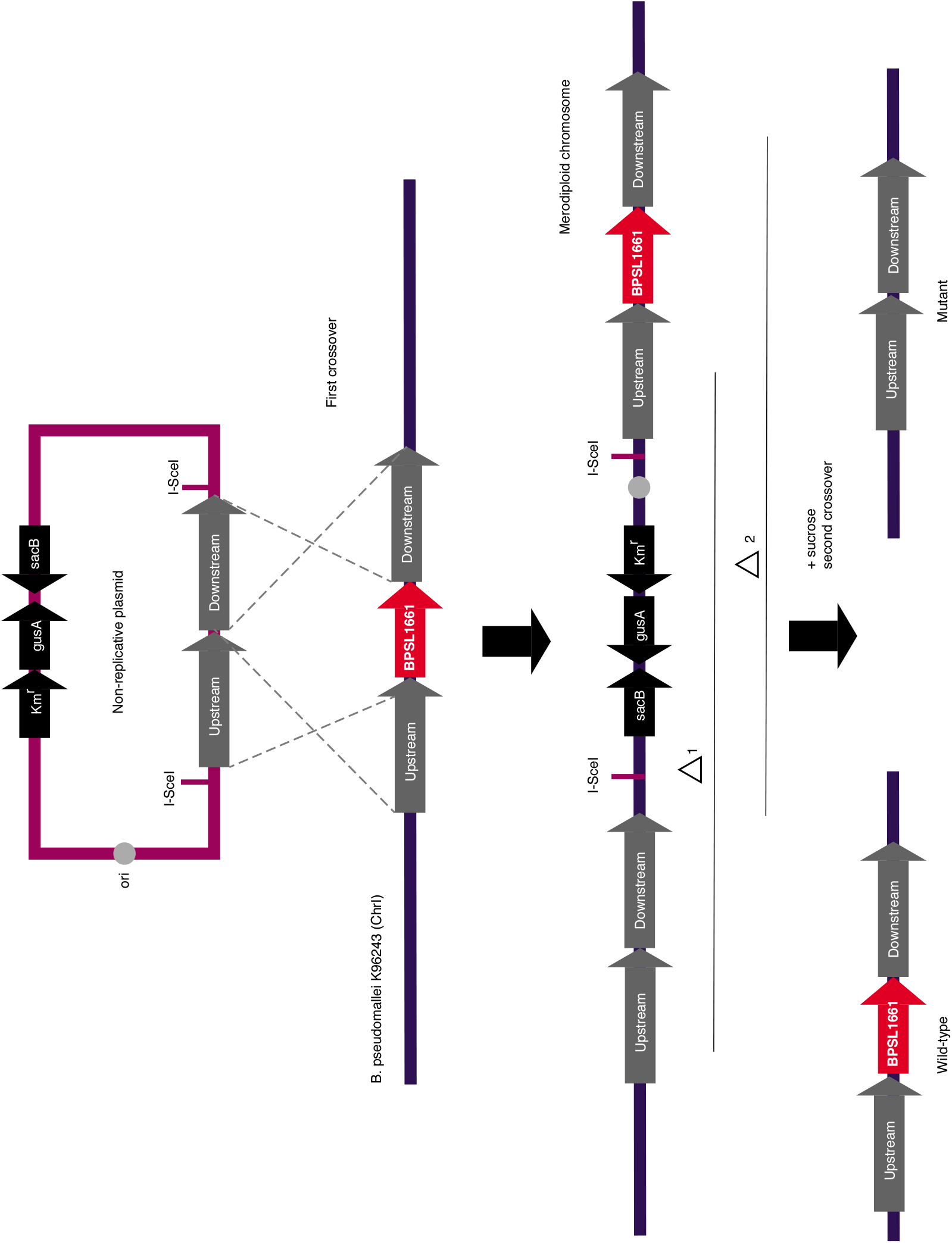
Construction of *BPSL1661* mutant. Schematic diagrams of plasmid-based allelic exchange of *B. pseudomallei* K96243 to construct *BPSL1661* mutant as modified from Lopez *et al*. 2009^49^. (a) a crossed PCR amplicon was generated from the assembled segments containing the upstream and downstream sequence of *BPSL1661* and cloned into pEXKM5, to construct the recombinant pEXKM5Δ*BPSL1661*;Km^r^. (b) The recombinant plasmids were integrated into chromosome of *B. pseudomallei* by homologous recombination (dash line denoted as the 1^st^ crossover). The integrated recombinant plasmid replaced the *BPSL1661* in the *B. pseudomallei* K96243 chromosome, called merodiploid. The sucrose counter selection was used to select for double crossovers (2^nd^ crossover), resulting in generation of either in a wild-type strain (Δ1) or *BPSL1661* mutant strain (Δ2). (A high resolution figure is also archived in https://figshare.com/s/df156bfe82883baf7efd)

